# Biophysical Essentials – A Full Stack Open-Source Software Framework for Conserved and Advanced Analysis of Patch-Clamp Recordings

**DOI:** 10.1101/2024.01.24.576881

**Authors:** David Zimmermann, Michaela Kress, Maximilian Zeidler

**Author notes:** **Corresponding authors:** Dipl.-Ing. David Zimmermann Maximilian Zeidler, PhD. Contributor Roles D.Z., M.K. and M.Z.: Conceptualization D.Z. and M.Z: Methodology; Validation; Formal analysis; Investigation D.Z. and M.Z.: Source Code Implementation; Data curation D.Z., M.K. and M.Z: Writing – original draft preparation; Writing – review & editing D.Z. : Visualization M.Z.: Supervision; Project administration.

## Abstract

Patch-Clamp recordings allow for in depth electrophysiological characterization of single cells, their general biophysical properties as well as characteristics of voltage- and ligand-gated ionic currents. Different acquisition modes, such as whole-cell patch-clamp recordings in the current or voltage clamp configuration, capacitance measurements or single channel recordings from cultured cells as well as acute brain slices are routinely performed for these purposes. Nevertheless, multipurpose transparent and adaptable software tools to perform reproducible state-of-the-art analysis of multiple experiment types and to manage larger sets of experimental data are currently unavailable. We therefore developed Biophysical Essentials (BPE) as an open-source python software for transparent and reproducible analysis of electrophysiological recordings. While initially designed to improve time consuming and repetitive analysis steps, BPE was optimized to cover entire workflows from data acquisition, preprocessing, visualization and normalization of single recordings up to stacked calculations and statistics of multiple experiments. BPE can operate with different file formats from different amplifier systems and producers. An in-process database logs all analysis steps for later review and serves as a central storage point for recordings. Statistical testing as well as advanced analysis functions like Boltzmann-fitting and dimensional reduction methods further support the researchers’ needs in projects involving electrophysiology techniques.

**Impact statement:** Biophysical Essentials provides fundamental support for transparent patch-clamp analysis to improve reproducibility in electrophysiology research.

## 1. Introduction

Since the first amplifiers were developed (Neher & Sakmann, 1976)the single electrode patch-clamp technique has developed into a widely used routine method in neuroscience and electrophysiology (Hamill et al., 1981). Starting from the characterization and pharmacology of nicotinic acetylcholine receptor, pharmacological interventions targeting specific ion channels including voltage gated Na^+^, K^+^ or Ca^2+^ channels, NMDA and AMPA glutamate receptors, GABA and glycine activated ion channels, and ryanodine-, inosital1,4,5-trisphosphate (IP3) or transient receptor potential (TRP) channel family members have been systematically examined with conventional patch-clamp recordings (Imming et al., 2006; Sakmann et al., 1980). The classical patch technique offers broad experimental flexibility and therefore represents the gold standard in basic research today. In order to speed up the pace in drug discovery and development, high throughput automated patch-clamp (APC) approaches are emerging, such as Flyscreen (Lepple-Wienhues et al., 2003), AutoPatch and RobotPatch (Vasilyev et al., 2005)or Synchropatch (Nanion Technologies GmbH) which allow to record from multiple cells at the same time for example by using a planar recording substrate instead of a classical patch electrode (Dunlop et al., 2008). Such high throughput experiments reveal valuable insights into biological variability in well-defined experimental flows (Seibertz et al., 2022).

Both, classical and APC technology, require detailed analyses of current and voltage traces to provide large amounts of high precision biophysical data. The analyses of these data requires significant expertise in signal processing, curve fitting and statistics which is supported by various existing commercial and open-source software tools that are optimised and tuned for specific amplifier systems and data formats such as Patchmaster and Fitmaster (current version PatchmasterNext (HEKA, n.d.) pClamp (current version is pClamp11 with Clampex, Clampfit and Clampfit Advanced) (*Friendly and Versatile Patch-Clamp Analysis with Axon PCLAMP 11*, n.d.) IgorPro (*Igor Pro from WaveMetrics | Igor Pro by WaveMetrics*, n.d.), OpenEphys (*Download the Open Ephys GUI — Open Ephys*, n.d.), Matlab (*Biological Sciences - Electrophysiology - MATLAB & Simulink*, n.d.), Neo (*Neo - NeuralEnsemble*, n.d.), Elephant (*Elephant - Electrophysiology Analysis Toolkit — Elephant 0.13.0 Documentation*, n.d.) and multiple more (*Free Alternatives to Commercial Electrophysiology Software*, n.d.)

We compared commonly used state-of-the art software tools within the context of typical neuroscience research scenarios together with analysis-relevant requirements and features of the respective software. In detail, these requirements compromise (1) *patch-clamp specific analysis functions*, (2) *extendable for new analysis functions*, (3) *transparency and reproducibility of analysis functions*, (4) *analysis of multiple cells/recordings in parallel*, (5) *analysis and result visualization according to cell-specific meta data*, (6) *cross-platform functionality* including Linux, Windows, MacOS and (7) *interoperability* between different patch-clamp result file formats. None of the existing software (listed and compared in Supplementary Table 1) addresses all of the identified requirements indicating that there is an unmet need for good-practice, transparent, universally applicable and reproducible data analysis of patch-clamp recordings’ data flows. To tackle this issue, we developed the Biophysical Essentials (BPE) full stack python software with an additional web interface to store, share and compare research data and findings. BPE allows for fast and conserved patch-clamp data analysis for multiple recordings simultaneously. It eliminates time consuming, error prone and repetitive analysis steps. The framework is contributed open-source to the entire scientific community and can be modified to meet research and lab specific needs individually.

## 2. Materials and Methods

### 2.1 Analysis of available software tools and identification of technical requirements

We investigated relevant criteria derived from two typical use cases in neuroscience and patch-clamp research. Scenario (A) resembled the characterization of electrophysiological excitability of cell populations e.g., with or without a specific gene knockout or mutation. Scenario (B) described research studying alterations in electrophysiological properties of a cell before and after the application of specific substance. Both scenarios would incorporate recordings of a cell’s response to various stimulation protocols, e.g. I-V relationship in the voltage-clamp configuration, rheobase determination, action potential (AP) fitting or a firing pattern analysis in the current clamp mode. This exemplary selection of stimulation protocols illustrated specific analysis needs with respect to a step or ramp protocol, normalization procedures dependent on the operation modes like voltage- or current clamp configuration, and fitting functions. These needs summarized as *patch-clamp specific analysis functions* criteria (1) were fully met by the three most referenced patch-clamp analysis tools Patch/Fitmaster, pCLAMP/ClampFit and IgorPro. Despite the broad heterogeneous analysis functions offered by these software tools, none of them satisfied criteria (2), *extendable for new analysis functions* like determination of a specific identified spiking pattern. In contrast, open source packages like Neo or the commercial programming and computing platform Matlab provided high degree of freedom options to append and integrate individual analysis functions into the data analysis process with existing/provided functions, however, require proficiency in programming as a prerequisite. Criteria (3) the *transparency and reproducibility of analysis functions* is satisfied by Patch/Fitmaster, pMaster/ClampFit and IgorPro for single files but not an entire file set with multiple recordings.

Addressing the introduced scenarios (A) and (B) would require a large number of repetitions. Therefore, the analysis of multiple cells with conserved parametrizations resembles a fundamental need in patch-clamp analysis which was summarized as criteria (4), *analysis of multiple cells in parallel*. The result presentation and the calculation of potential mean traces requires appropriate handling of the meta data of the cells. This need is resembled by criteria (5), *analysis and result visualization according to cell-specific meta data*. None of the tools Patch/Fitmaster, pMaster/ClampFit or IgorPro satisfied criteria (4) and (5). To release electrophysiology research from technical limitations and to improve reproducibility and transparency, we added criteria (6), *cross-platform functionality* including Linux, Windows, MacOS and criteria (7), *interoperability between different file formats*, such as .dat and .abf files and open file formats such as neurodata without borders (Teeters et al., 2015) Patch/Fitmaster, pMaster/ClampFit or IgorPro available for Windows and MacOs and Patch/Fitmaster allow data export into an IgorPro supported file format. Igor Pro itself supports multiple file formats including HDF and HDF5, Matlab files and multiple more (Supplementary Table 1).

### 2.2 BPE packages and hardware requirements

The source code of BPE was implemented in Python (current version 3.11) and uses PySide6 (version 6.5.1.1) as the official Python module from the QT for Python project, which provided complete access to the QT 6.0+ framework. Button images were selected from ICONS8 (*Icons8*, n.d.). Graphics were generated using the python packages *Matplotlib*, *PyQtGraph*, and *Seaborn*. All listed software packages were selected to enable BPE execution on Linux (Ubuntu, 23.04), MacOS (macOS 13) and Windows (Windows 10/11) as well as on x86 and ARM architecture. Generally, the minimum hardware requirements include 4 GB RAM and a 2.5 GHz CPU. Memory usage depends on the amount and the size of data stored in the local database. The import of data from the recording files into the database as well as the application of analysis functions support multithreading to speed up the most time-consuming procedures. Current package versions were provided within the respective .yml file in the github repository https://github.com/ZiDa20/Biophysical_Essentials.

### 2.3 Amplifier Communication and Experimental Setup

For BPE communication with the HEKA amplifier (EPC 10), a valid version of HEKAs control software Patchmaster (tested with Patchmaster v2x90) is required via the batch communication (BCOM) feature. BCOM allows Patchmaster to read and execute control commands from an “in-out file” that we manipulate with BPE by updating the command id and the command to be executed. The entire list of available batch communication commands can be found in HEKAs documentation.

### 2.4 Data Import and local Database Management System

Either live recorded data or previously acquired recordings can be imported into BPE framework from HEKA amplifiers (.dat file format) and amplifiers from Molecular Devices (*.abf file format). To read *.dat files we used a modified version of the publicly available heka_reader package with pgf file reading support (github campagnola/heka reader) adapted from the official matlab heka dat reader. To enable *.abf support, the pyabf-package was used (*PyABF - A Simple Python Library for Working with ABF Files*, n.d.). Files are imported into an in-process SQL online analytical processing (OLAP) database management system based on DuckDB (Raasveldt et al., 2019)because of its high-performance design to process large amounts of data. BPEs offline analysis module operates on the permanent and locally stored database while during online analysis, files are imported into an in-memory database. As one measure to keep the permanent database at a minimum dataspace level and to avoid sources of inconsistency it is not accessible after the restart of the software. Recordings viewed in the online analysis module can be transferred into the permanent local database. Further additions in terms of metadata and image data can be added to the database.

### 2.5 Statistics

The statistics module compromises t-test, Welchs-test, Mann-Whitney-U test, paired-t-test, Wilcoxon-signed-rank-test, Kruskal-Wallis test and ANOVA. Therefore, result data distribution is automatically tested using Shapiro-Wilk test, data variance is tested using Levene-test. The appropriate statistical test will be suggested according to the number of specified meta data groups, data distribution, variance and data dependency. All the tests were performed using *scipy stats* package.

### 2.6 Patch-Clamp Recordings

DRG neurons were harvested as described [PMID 34252913], plated and incubated in culture medium at 37°C and 5% CO_2_ for 24 hours. Neurons derived from induced pluripotent stem cells were differentiated as described in [PMID 34486248]. Whole cell patch-clamp recordings were performed using a HEKA EPC 10 amplifier. The cells were kept in extracellular solution (ECS) containing NaCl (145 mM), HEPES (10 mM), D-Glucose (10 mM), KCl (5 mM), CaCl_2_ (2 mM) and MgCl_2_ (1 mM) at a pH of 7.3 adjusted with NaOH. Intracellular solution (ICS) was used to fill borosilicate glass pipettes pulled with a horizontal puller (P-1000, Sutter Instrument Company), with a pipette resistance between 3-5 MΩ. The ICS contained potassium D-gluconate (98 mM), EGTA (5 mM), HEPES (10 mM), KCL (45 mM), MgCl_2_ (2 mM), Na_2_GTP (0.2 mM), MgATP (2 mM) and CaCl_2_ (0.5 mM), with a pH adjusted to 7.3 using KOH.

## 4. Results

Biophysical Essentials compromises five modules: an experimenter module, an online analysis module, an offline analysis module, a database viewer module and a BPE online service module. These modules function independently from each other but can be also concatenated to an analysis pipeline depending on individual demands as illustrated in Figure 1A.

**Figure 1:**
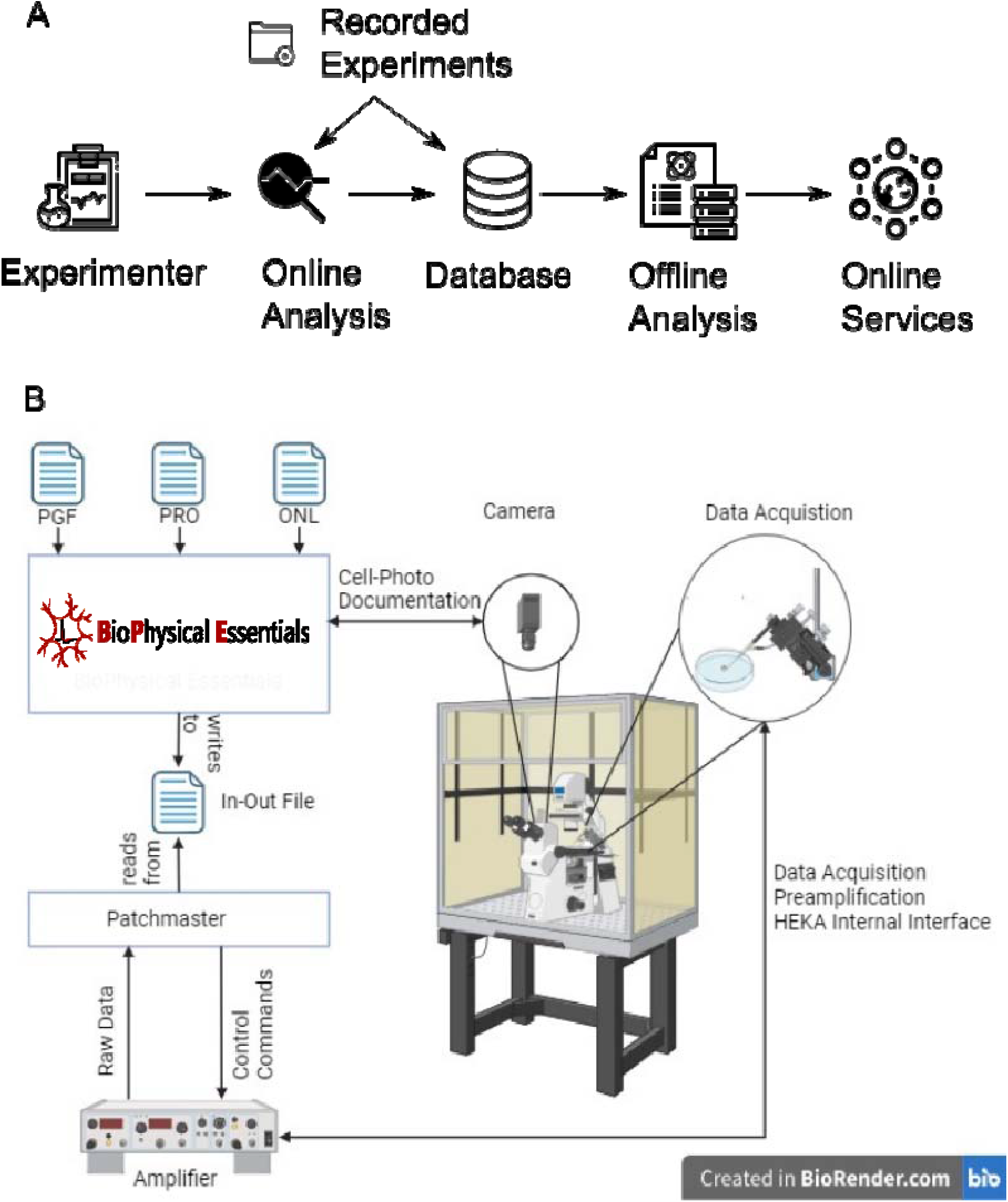
Modular Composition of BPE. A) Five different modules allow BPE to provide a full stack analysis pipeline represented by independently functioning modules experimenter, online analysis, offline analysis, database and online analysis. B) The experimenter module requires the input of a pulse generator file (.pgf), a protocol file (.pro) and an online analysis (.onl) file generated by Patchmaster. To establish a batch communication, Patchmaster needs be set as a receiver and will therefore search for a specific in-out-file in a determined directory. BPE writes commands to this in-out-file according to HEKA’s provided documentation of the batch communication interface. Created with BioRender.com

### 4.1 Experimenter Module

BPE can be integrated into the patch-clamp experiment workflow from the very beginning of the data collection, starting at raw data acquisition. Currently, BPE supports communication with the HEKA EPC 10 amplifier and requires a running Patchmaster software with enabled batch communication (details are described in the methods section). The experimenter module (EM) was designed to select and execute electrical stimulation protocols, establish the hardware connection with the amplifier and an optional camera device (Figure 1B). EM offers to add recording-relevant additional information, such as the composition of intra-, extra- and optional stimulation solution. For manual parametrization of amplifier input, such as adjusting the injected current or voltage within a protocol, execution of protocols via Patchmaster remains crucial. During raw data acquisition, EM operates in parallel to the existing hardware and software pipeline from HEKA, by fetching the recorded data from the Patchmaster notebook via the opened batch connection. If the EM executing computer is connected with a camera attached to the microscope, photo documentation of the patched cells can be activated. Once the execution of a stimulation protocol is finished, the recorded data, as well as the applied protocol, are displayed in BPE’s online analysis module. From here, the recorded data can be imported into the BPE local database while the original recording file remains unaltered. All provided metadata information of the experiment are added to the database too. This allows for in-depth metadata analysis, batch correction as well as normalization methods in later analysis steps as we will show in the offline analysis module.

### 4.2 Online Analysis Module

The online analysis (OA) module is designed to serve as a viewer module of individual recording traces, provides a digital lab book to enter additional experiment information and allows to transfer single traces into the BPEs database (described in the next section). In BPE, recordings are generally interpreted as a structure with three levels: experiment, series, and sweeps. Experiments refer to a single recorded cell, series represent a specific stimulation/recording protocol that was performed on this cell, and sweeps represent individual voltage or current steps of the individual stimulation protocols. Within the graphical user interface (GUI), stimulation and recording data are displayed simultaneously which allows a good overview and evaluation of the cell specific response to changes in current, voltage or other stimulation types. The digital lab book allows for manipulation of existing metadata or entering new data and comments in a table-like structure. If combined with the EM, intra- and extracellular solutions, pipette resistance, performed compensation, and additional required parameters or comments are transferred from the EM into the lab book automatically. Also, recorded videos and images of the cell throughout the experiment are appended to the lab book. Recordings, their annotated meta data and the digital lab book can be transferred from the OA module into BPE’s local database to become available for further data analysis.

### 4.3 Database Module

BPE stores all data and analysis steps and details within an online analytical processing (OLAP) database (DB) that is only stored locally and always operates in the background as soon as BPE is started. The main structure of the distinguishes between raw data and analysis data, as indicated in the entity-relationship diagram in Figure 2. To keep the database as small as possible, all experiments are constrained to be unique in the database by the experiment name and a series identifier. Via mapping tables, an experiment can be assigned to multiple different analyses and each analysis is identified by a unique identifier. The entire offline analysis process, which will be described in the next section, is stored in the database and can be reconstructed at any later time point from the database entries. The database content is represented within a dashboard as shown in Supplementary Figure S3. Performed analysis within BPE can be exported from one computer and imported onto another computer via implemented import and export functions.

**Figure 2:**
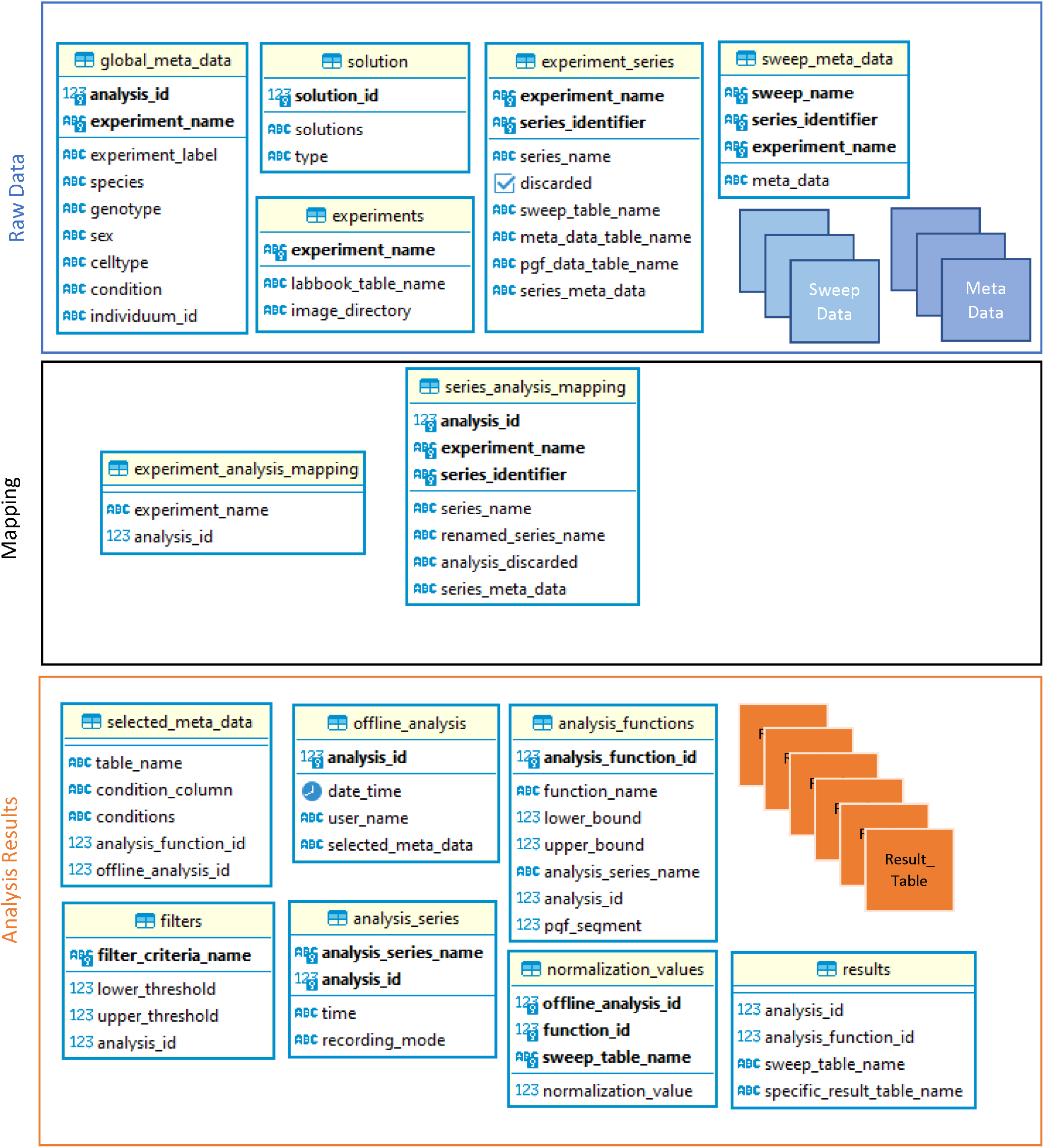
Entity Relationship Diagram of the local database structure: BPE organizes its internal data structure within a local database based on the DuckDB architecture. Raw data must be imported once into the DB, the recording file itself will be never modified. All further processes are performed on the database data only. To keep the size of the database small, raw recording data are unique within the DB while they can be mapped to multiple analysis via the introduced mapping tables.

### 4.4 Offline Analysis Module

The core of BPE is its offline analysis (OFA) module, which enables the analysis of multiple patch-clamp recordings in parallel according to annotated meta data. The analysis process is composed of three steps, which compromise (1) (manual) data control, (2) analysis setup and (3) result visualization. Once performed, analysis can be reopened from the database quickly. To start a new independent analysis, new recordings can be imported from the original recording files into the database as well as already imported recordings can be selected from the DB via the DB dashboard. The analysis was designed to cover comparisons between different cells or within the same cells. For SQL-experienced users, advanced data selection options are available.

The first panel of the OA shows the structural composition of each single cell and performed protocols within in a hierarchical tree view as shown Figure S2. The structural view is supported by a graphical visualization for cell specific stimulation protocols and the electrical responses. This aggregated and comprehensive data assembly and graphical visualization allows an easy and manual control of signal traces and electrical properties. Experiments and series that do not fulfill quality criteria, such as low leakage or noise levels, can be manually removed from the analysis. Graphical visualization and individual data selection allow OFA to be applied in different biological context, such as peripheral neurons, pancreas cells or myocardial cells. The manual filtering process can be extended and mimicked by automatic filter options, which allow thresholding of relevant parameters, such as the capacitance of the cell (as a measure of the cell size) or serial resistance (as a measure of the series resistance between the pipette and the cell) throughout the experiment. Experiments can be further separated according to annotated metadata, such as genotype, sex, or condition. This first panel of the OFA includes manual data screening and filtering of the input data to gain a maximum of control over performed patch-clamp-recordings and the downstream analysis. In the second panel, analysis functions can be selected and parametrized related to the respective protocol, e.g. analysis for all IV recordings or analysis for all rheobase recordings respectively. Therefore, OFA provides various default and protocol-specific 1^st^ level analysis functions listed in Table 2. Multiple independent functions can be applied to one specific protocol. Also, dependent analysis functions, such as subtracting the maximum value from interval *x* and the mean value from an interval *y*. The respective intervals can be selected by movable cursor bounds. For voltage-clamp protocols, normalization of the recorded currents to the cell membrane size (CSLOW-parameter) (Ismaili et al., 2020) or a manual normalization parameter are supported. Both methods are implemented in OFA and can be selected from the front end.

Once the analysis was set up respective to the specific protocol and data quality, existing meta data groups, normalization method and analysis functions, the results will be determined, stored in the DB and will be displayed immediately. In the graphical result representation, outliers can be selected to go back to the raw traces and study possible reasons. In addition, numerical output of the results is provided for each analysis function. Various 2^nd^ level functions can be selected for further analyses like dimensional reduction und cell clustering or IV curve fitting. For quantitative assessment of differences between experimental groups, a statistics module was implemented and provides statistical test to be applied to the generated results. The number of the comparison groups, data distribution (via Wilcoxon-Rank-Sum test) and variance (via Levene-test) are accessed automatically by OFA and suggest appropriate statistical tests. However, tests can be also selected manually. The full documentation of BPEs workflows with additional videos and further descriptions is available at https://biophysical-essentials.i-med.ac.at/.

### 4.5 Validation

To ensure correctness of the implemented analysis functions and workflows in BPE, all 1^st^ order analysis functions listed in table 1 were carried out on a dataset of ten cells using Patchmaster v2x90.5 “manually”. Numerical values were compared individually for each recording and analysis function as summarized in Supplementary Table 3. To ensure that BPEs meta data handling and meta data specific workflows result in the same values as manual meta data specific analysis, the recordings were separated into two arbitrary groups of five cells. The mean values of the respective groups were compared for each analysis function as described before. Eventually, to validate proper function of the normalization algorithms, additional analysis with and without normalization to the C-slow parameter was carried out. As demonstrated in Supplementary Table 3, manual as well as BPE performed calculations resulted in the same numerical values for not normalized, normalized and meta data grouped results. To ensure that further implementations and adoptions to BPE will not interfere with the validated result calculations, these validations will be performed prior to every update to the main code.

**Table 1:** Summary of implemented analysis functions and brief description.

|  | Analysis Function | Description |
| --- | --- | --- |
| 1<sup>st</sup> order | Max, Min, Mean Amplitude | Within a given interval, determination of maximum- or minimum amplitude, or calculation of the mean amplitude within the interval |
|  | Time to Max, Min | Time to the maximum or minimum of a signal |
|  | Action Potential Fitting | Determination and calculation of action potential characteristics, e.g. threshold, maximum, half width and 15 other parameters (Supplementary Table 2) |
|  | Rheobase Detection | Detection of the minimum current required to initiate an action potential |
|  | Peak Detection | Detection of peaks |
|  | Input Resistance | Linear regression to determine the input resistance of a cell |
|  | Rheoramp-Detection | Determination of action potentials per sweep |
|  | Firing Pattern Analysis | Analysis of the spike pattern considering AP count, frequency and bursting, tonic or phasic-properties |
| 2<sup>nd</sup> order | PCA | Linear dimensional reduction method: Principle Component Analysis |
|  | IV-curve Fitting | Boltzmann fit |
|  | Hill-Function-Fitting | Determination of ligand-receptor binding relationship |

### 4.6. Classification and characterization of neurons using BPE

Translating knowledge and research findings from animal into human model systems represents a major task in neuroscience and medical research. Nociceptors derived from induced pluripotent stem cells (iNocs) are emerging as promising preclinical model to study human nociception (Schoepf et al., 2020; Zeidler et al., 2021). Since BPEs analysis functions are not restricted to specific biological context, it can be used to process patch-clamp recordings from both, iNocs as well as murine sensory neurons obtained from dorsal root ganglia (DRG). As a first experiment to document BPE’s versatile functionalities, DRG neurons’ and iNocs’ specific electrical characteristics were accessed by conducting patch-clamp recordings and action potential (AP) properties were compared (Figure 3A-D). Following the workflow illustrated in Figure 1A, raw data were imported from the recording file into the DB of BPE with the respective meta data annotation (iNoc or DRG), a new offline analysis was started and action potential fitting, peak detection analysis and PCA were performed for five action potentials (n=9, Figure 3A). Characteristic AP mean traces for iNocs and DRG neurons were determined and aligned along the AP-maximum to qualitatively compare the mean AP signal shapes. In Figure 3B, representative mean traces are shown within a 10ms window before and after the AP maximum. For quantitative comparison, AP-fitting was performed for 18 parameters of interest and for each recorded cell individually, followed by mean value determination (Figure 3C, Supplementary Table 2). Based on the 18 AP-fitting parameters, principal component analysis (PCA) was performed and revealed a dimensional separation between iNocs and DRG neurons (Figure 3D). The BPEs statistics module was used to identify statistical differences between the AP characteristics of the two cell populations. Independent t-test analysis between AP parameters of DRG neurons and iNocs revealed significant differences between most AP parameters. Detailed mean values, standard deviation and statistical significance are summarized in Supplementary Table 2.

**Figure 3.**
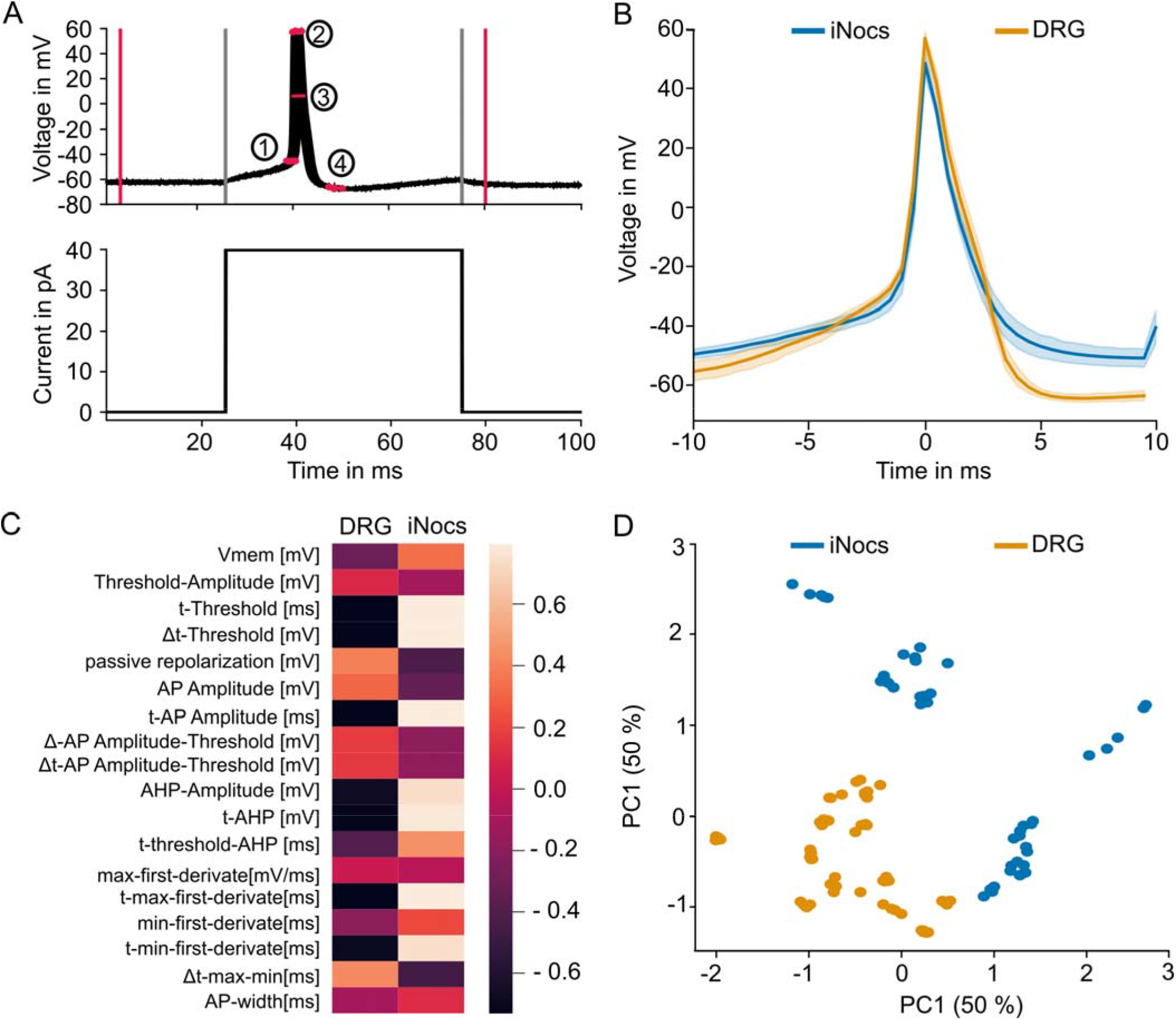
Comparison of Action potential characteristics of mouse DRG neurons and human iNocs. (A) Representative stimulation protocol and the recorded action potential of the cell. The protocol was repeated 5 times. The red vertical lines indicate the analysis interval, the light grey horizontal lines separate the 3 intervals defined in the pulse generation file. Markers 1-4 illustrate the result of the enabled live analysis feature. For action potential fitting, V_thresh_ (1), V_max_ (2), t_half width_ (3) and V_AHP_ (4) are displayed as subsitutes. (B) Comparison of the mean trajectory between 9 iNocs and 9 DRG neurons. Detected action potentials were aligned by their maximum peak V_max_. (C) Action potential fitting was performed for 18 parameters and the mean for each cell type was scaled and visualized. Mean data and standard deviation are summarized in supplementary table 2. (D) PCA was applied to the AP fitting parameters of each single AP and principal components 1 and 2 were plotted accordingly.

## 5. Discussion

In this work, we have implemented and introduced BPE, a new full stack Python software tool which aims to compensate for lacking features in current state of the art patch-clamp recording analysis software (Supplementary Table 1). BPE was designed to automate repetitive and error-prone patch-clamp analysis steps while offering a maximum of manual control of recording quality and data specificity through the implemented GUI, as well as general and patch-clamp specific analysis functions without limitation to the biological context. The complex offline analysis module addresses the needs for patch-clamp specific analysis functions (criteria 1, Table 1), is extendable for new analysis functions (criteria 2), provides different analysis functions for multiple cells simultaneously (criteria 3) and considers cells meta data within the analysis (criteria 4). In addition to basic analysis functions like the detection of the minima, maxima, means or respective time to specific extrema, BPE provides specific functions for action potential fitting, determination of the input resistance or membrane potential, rheobase detection and analysis, firing pattern analysis as well as Boltzmann- and Hill-fittings which resemble analysis functions regularly applied to experiments assessing neuron excitability (Madden et al., 2020; Weth-Malsch et al., 2016). BPE provides additional analysis functions such as dimensional reduction analysis to study overall similarities and consistency between cells or experiments.

To document BPE’s applicability for reliable and rapid analysis of patch-clamp recordings and performance of respective experiments, we have compared electrical AP characteristics between mouse and from human pluripotent stem cell derived nociceptors (iNocs) (Figure 3), as introduced in various translational pain research studies (Alsaloum & Waxman, 2022; Zeidler et al., 2021; Zurek et al., 2023). Dimensional reduction analysis using PCA revealed the highest similarities of AP characteristics within signals from the same neuron but also within the respective iNoc or DRG neuron category (Figure 3D). However, the parameter-specific analysis revealed significant differences for most of the AP parameters between human and mouse sensory neurons. Integration of our own data with other patch-clamp data sets e.g. from human primary sensory neurons (Tiwari et al., 2023; Zurek et al., 2023) in BPE offers great potential to elucidate functional species dependent differences which are already demonstrated at the RNA and protein level (Ray et al., 2018). To enable build up of comprehensive electrophysiological data sets, data reanalysis and maximum exploitation of already available and new data, an online platform generated within the BPE environment with the community, independent from the patch clamp amplifier system, has been launched at https://biophysical-essentials.i-med.ac.at/. This platform aims to foster reproducibility, transparency and enable comparison of data from different laboratories, species and model systems. BPEs ability to keep track of intra-and extracellular recording solution as well as stimulation protocols enables to compare data taking into account different recording conditions such as ionic strength and will be further developed to correct for these differences relevant for data comparison. BPE extends beyond already available patch clamp specific, open source and full stack analysis tools such as the python tool PatchView (Hu & Jiang, 2022) or the C++ based software StimFit (Guzman et al., 2014). since it was developed in particular for reproducible analysis workflow which is documented in detail and stored in the background database. The entire workflow ranging from raw data selection and filtering to analysis function parametrization and result visualization of any already performed analysis can be reviewed at any time which provides genuinely new value in terms of workflow transparency.

BPE is distributed under open-source license (criteria 5) and thus allows to join forces with existing implementations to further improve BPE’s analysis function stack (Table 1) and custom analysis function development. The cross-platform operability is covered by integration tests for Ubuntu (tested with 22.04 LTS), MacOs (tested with 12) and Windows (tested with Windows 11®) (criteria 6). In comparison to pure data analysis tools like IgorPro or PatchView, BPE additionally provides interaction with amplifier- and camera hardware thus allowing for process automatization and collection of other data types, such as a image documentation of the recorded cell. Integration and analysis of images will allow to consider general cell morphology parameters for correlation with electrophysiological parameters which will generate novel insight into the correlation of functional and morphological characteristics.

## 6. Future Outlook

BPE integrates and processes data from available databases, analysis functions and graphical visualizations at high veleocity by making optimal use of available hardware resources. Currently, the HEKA (.dat) as well as the AXON file formats are supported. Based on its versatile philosophy, BPE is suitable for implementing new datatypes, including data emerging from high throughput or automated recording systems such as the Syncropatch environment. Also, BPE will be equipped with more complex statistical tools like generalized linear mixed model and linear regression model. New analysis functions such as ePSC/iPSC signal detection as well as additional normalization methods, such as t-normalization will further improve BPE’s function stack, (Golowasch et al., 2009)

## 7. Data Availability

The source code of BPE is available on Github: https://github.com/ZiDa20/Biophysical_Essentials. The website for documentation, and online data access or sharing is accessible via https://biophysical-essentials.i-med.ac.at/

## 8. Conflict of Interest

No conflicts of interest have to be declared.

## 9 Acknowledgement

We are particularly grateful to Michael Hörtnagl and the IT-Department of Medical University Innsbruck for providing expert support in deploying the projects website on the local webserver.

This work was founded by the Austrian Science Fund FWF I-050870.

---

**Supplementary Figure S1:** Data Visualization of Available Patch Clamp Data

**Supplementary Figure S2:** Offline Analysis Panel 1: Quality Control

**Supplementary Figure S3:** Database Viewer: Data availability

**Supplementary Figure S4 :** Offline Analysis Panel 2: Analysis Function Setup

**Supplementary Figure S5:** Offline Analysis Panel 3: Result Visualizer: Graphical Representation

**Supplementary Figure S6:** Offline Analysis Panel 3: Result Visualizer Statistics Module

## Notes

### Competing Interest Statement

The authors have declared no competing interest.

