## Supplementary Figures S1-S6 for "Biophysical Essentials – A Full Stack Open-Source Software Framework for Conserved and Advanced Analysis of Patch-Clamp Recordings"

### Slide 1
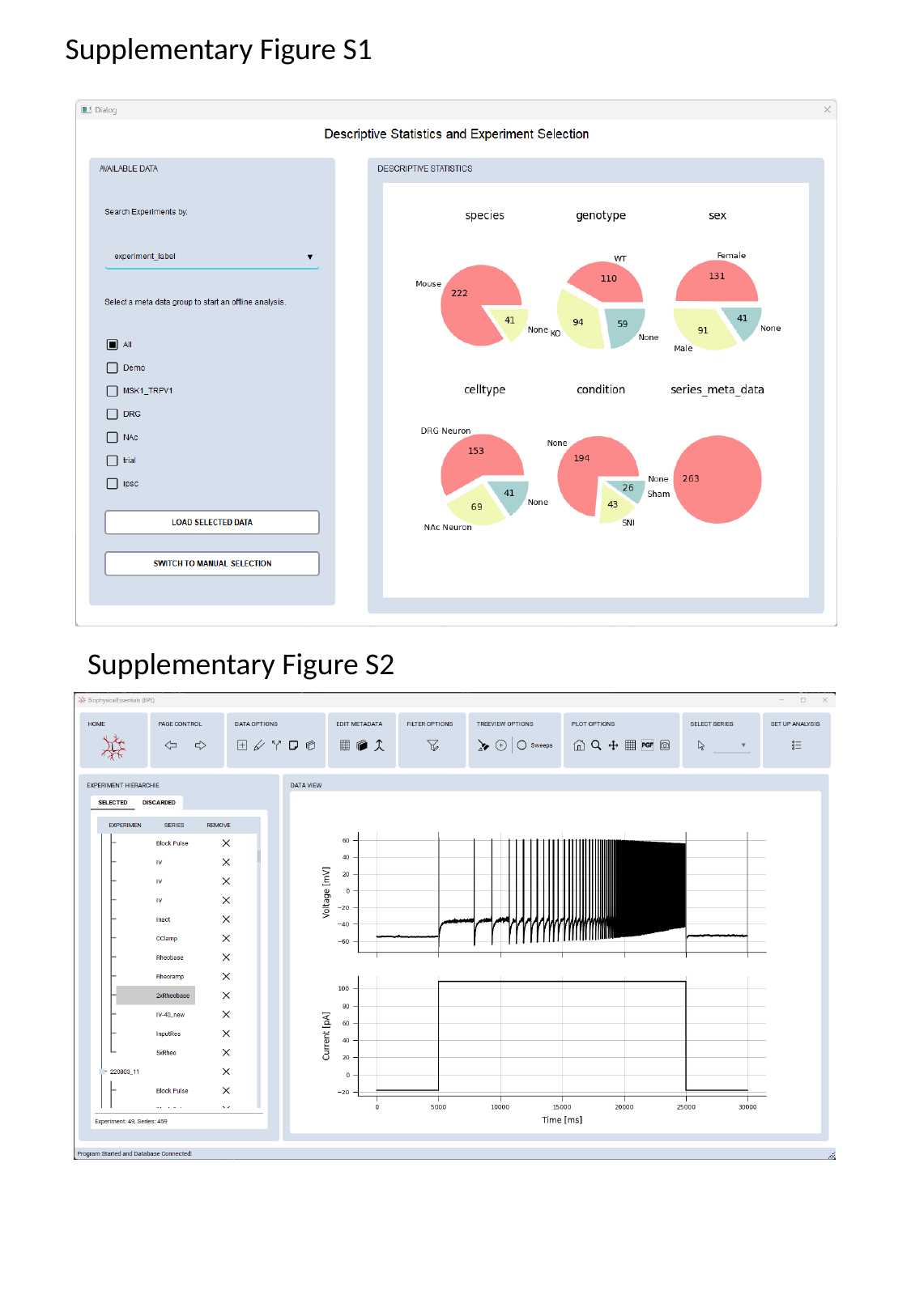

Supplementary Figure S1
Supplementary Figure S2

### Slide 2
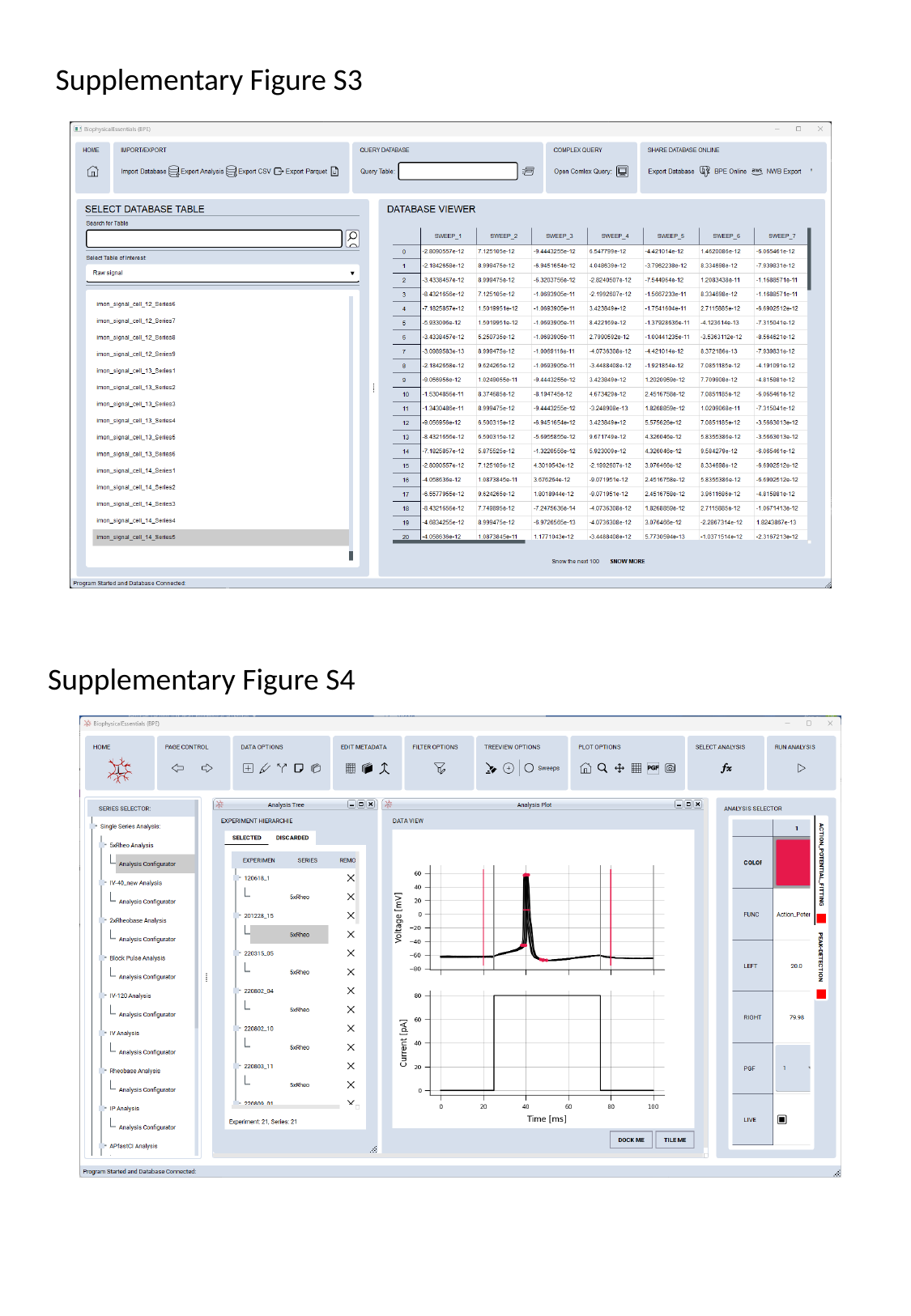

Supplementary Figure S3
Supplementary Figure S4

### Slide 3
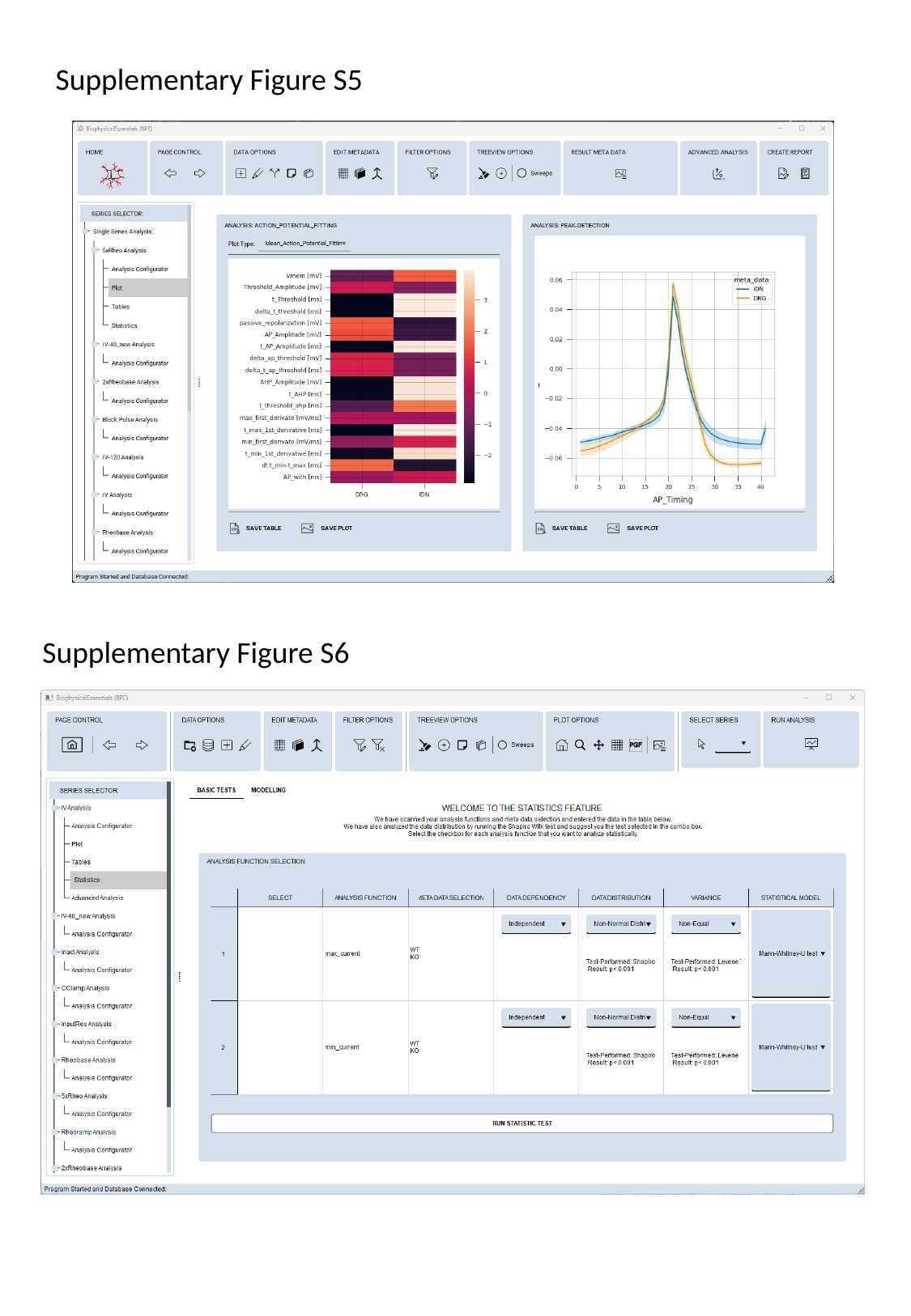

Supplementary Figure S5
Supplementary Figure S6
