## Supplementary Tables 1-2 for "Biophysical Essentials – A Full Stack Open-Source Software Framework for Conserved and Advanced Analysis of Patch-Clamp Recordings"

**Supplementary Table 1: Comparison between State of the Art patch-clamp analysis software Patchmaster/Fitmaster, pCLAMP, IgorPro and BPE**

| **Criteria** | **Description** | **Patch/ Fit**  **master (Next)**  **HEKA** | **pClamp**  **Molecular Devices** | **Igor Pro** | **BioPhysical Essentials** |
| --- | --- | --- | --- | --- | --- |
| **1** | **Patch-Clamp Specific Analysis Functions** | Action Potential Fitting  Event Detection | Fittings  Event Detection  Population Spike Search | Curve Fitting  Peak Detection | AP-Fitting  IV-Fitting  Hill-Curve-Fitting  Rheobase  Input Resistance  Firing Pattern  Capacity Measurements |
| **2** | **Extendable for new Analysis** | No | No | Yes | Yes |
| **3** | **Transparency and reproducibility of analysis functions** | Single files | Single files | Single files | Single and Multiple Files |
| **4** | **Analysis of multiple cells/recordings in parallel** | No | No | No | Yes |
| **5** | **Analysis and result visualization according to cell-specific meta data** | No | No | No | Yes |
| **6** | **Cross Platform Operability** | Windows,  (MacOS) | Windows,  MacOS | Windows,  MacOS | Windows,  MacOS, Linux |
| **7** | **Data Format Interoperability** | .dat | .abf | Delimited text, Fixed-field (FORTRAN) text, General Binary, Excel spreadsheet, HDF, HDF5, Matlab, JCAMP, Nicolet Instruments, SDTS DEM and DLG, National Instruments TDM (DIAdem) | .dat, .abf |

**Supplementary Table 2: Comparison of specific action potential characteristics between mouse dorsal root ganglia neurons and human induced nociceptors.**

|  | **Parameter** | **Unit** | **DRG** | | **iNocs** | | **p-value** |
| --- | --- | --- | --- | --- | --- | --- | --- |
|  |  |  | **mean** | **SD** | **Mean** | **SD** | **Independent t-test** |
| 1 | V_mem_ | mV | -60.76 | ± 4.10 | -56.21 | ± 2.79 | 0.002 |
| 2 | V_thresh_ | mV | -30.46 | ± 4.95 | -32.87 | ± 4.25 | 0.197 |
| 3 | t_thresh_ | ms | 34.41 | ± 2.98 | 51.08 | ± 4.90 | < 0.001 |
| 4 | Δt_thresh_ | ms | 9.41 | ± 2.98 | 26.08 | ± 4.90 | < 0.001 |
| 5 | V_pass. depolarization_ | mV | 30.30 | ± 5.16 | 23.34 | ± 2.15 | <0.001 |
| 6 | V_max_ | mV | 57.15 | ± 4.55 | 48.75 | ± 7.14 | <0.001 |
| 7 | t_max_ | ms | 36.14 | ± 2.89 | 52.62 | ± 5.01 | <0.001 |
| 8 | ΔV_max-thresh_ | mV | 87.61 | ± 5.45 | 81.62 | ± 10.86 | 0.0703 |
| 9 | Δt_max-thresh_ | mV | 1.73 | ± 0.36 | 1.53 | ± 0.18 | 0.0974 |
| 10 | V_AHP_ | mV | -65.51 | ± 2.01 | -51.18 | ± 4.98 | <0.001 |
| 11 | t_AHP_ | ms | 41.75 | ± 3.51 | 60.04 | ± 5.74 | <0.001 |
| 12 | Δt_AHP-thresh_ | mV | 7.34 | ± 0.89 | 8.95 | ± 1.20 | <0.001 |
| 13 | dV/dt_max_ | mV/ms | 133.94 | ± 11.96 | 131.40 | ± 29.39 | 0.7648 |
| 14 | t(dV/dt_max_) | ms | 35.49 | ± 2.87 | 51.87 | ± 4.95 | <0.001 |
| 15 | dV/dt_min_ | mV/ms | -55.63 | ± 7.00 | -50.71 | ± 12.74 | 0.2135 |
| 16 | t(dV/dt_min_) | ms | 38.01 | ± 3.25 | 53.27 | ± 5.14 | <0.001 |
| 17 | Δt (dV/dt_max-min_) | ms | 2.52 | ± 0.70 | 1.41 | ± 0.33 | <0.001 |
| 18 | t_half width_ | ms | 1.72 | ± 0.24 | 1.82 | ± 0.39 | 0.411 |
